# Sexual imprinting and speciation in two *Peromyscus* species

**DOI:** 10.1101/145243

**Authors:** E.K. Delaney, H.E. Hoekstra

## Abstract

Sexual isolation, a reproductive barrier, can prevent interbreeding between diverging populations or species. Sexual isolation can have a clear genetic basis; however, it may also result from learned mate preferences that form via sexual imprinting. Here, we demonstrate that two sympatric species of mice—the white-footed mouse (*Peromyscus leucopus*) and its sister species, the cotton mouse (*P. gossypinus*)—hybridize only rarely in the wild despite co-occurrence in the same habitat and lack of any measurable intrinsic postzygotic barriers in laboratory crosses. We present evidence that strong conspecific mating preferences in each species result in significant sexual isolation. We find that these preferences are learned in at least one species: *P. gossypinus* sexually imprints on its parents, but in *P. leucopus,* additional factors influence mating preferences. Our study demonstrates that sexual imprinting contributes to reproductive isolation that reduces hybridization between otherwise interfertile species, supporting the role for learning in mammalian speciation.

## INTRODUCTION

Sexual isolation, where sexual interactions such as divergent mating preferences or courtship behaviors reduce interbreeding, is a prevalent premating reproductive barrier that may facilitate speciation. Relative to some intrinsic postzygotic reproductive barriers, sexual isolation can accumulate rapidly among young allopatric (e.g. Mendelson 2003) and sympatric species (e.g. Coyne and Orr 1989, 1997), and it often acts as a major reproductive barrier among incipient sympatric species pairs (Coyne and Orr 1997; Noor 1997; Ramsey et al. 2003; Boughman et al. 2005; Nosil 2007; Matsubayashi and Katakura 2009). In several cases, sexual isolation is the sole reproductive barrier preventing hybridization between sympatric species, indicating that sexual isolation alone can be strong enough to reduce hybridization and thereby maintain genetic differentiation (e.g. Seehausen 1997; Fisher et al. 2006). Yet, despite the role that sexual isolation can play in instigating or maintaining reproductive isolation among species, its mechanistic basis—whether mating preference is genetic or learned—is often unknown.

Sexual isolation can evolve when mating traits and preferences are genetically encoded. If polymorphisms exist at a mating-trait locus and a preference locus, divergent alleles can co-evolve and fix between a pair of populations causing assortative mating. This scenario is known as a “two-allele mechanism” of reproductive isolation because two alleles must be present at both the mating-trait and preference loci (Felsenstein 1981). With the exception of a single pleiotropic trait/preference locus (Smadja and Butlin 2011), sexual isolation formed by the two-allele mechanism will break down due to recombination between the separate trait and preference loci unless strong selection, weak gene flow, or a high degree of linkage disequilibrium exists (Felsenstein 1981).

Sexual isolation can also evolve without genetically encoded preferences. Under a “one-allele mechanism” of reproductive isolation, a single allele yields assortative mating—for example, because of self-referent matching, mechanical assortment, or philopatry (Kopp et al. *in press*). Sexual imprinting, a process in which offspring learn to prefer familial traits at a young age (i.e. those of a mother, father, or siblings), has been considered an “one-allele mechanism” (Verzijden et al. 2012a) because populations that diverge in a sexually imprinted mating trait can mate assortatively thus leading to sexual isolation. Mechanisms such as sexual imprinting are arguably more efficient at establishing reproductive isolation than the above-mentioned two-allele mechanisms because they are immune to genetic recombination: separate preference alleles do not need to be associated with polymorphisms in mating-trait alleles to produce assortative mating (Felsenstein 1981; Smadja and Butlin 2011). Moreover, several theoretical models have shown that learned mating preferences will maintain sexual isolation much longer in populations experiencing gene flow than if mating preferences had a genetic basis because sexual imprinting lowers the amount of divergent natural selection needed to isolate groups (Laland 1994; Verzijden et al. 2005). Sexual imprinting may also boost reproductive isolation through reinforcement (Servedio et al. 2009) or by driving divergence in mating traits. If offspring develop preferences for more extreme versions of the traits on which they have sexually imprinted, peak shift can occur (ten Cate and Rowe 2007), which can in turn drive mating-trait evolution (ten Cate et al. 2006) and promote adaptive radiation (Gilman and Kozak 2015).

While sexual imprinting has long been recognized as a phenomenon that occurs within species, its potential impact on speciation has become better appreciated only over the last two decades (Irwin and Price 1999). It is a phenomenon that occurs in species with parental care, and has now been documented in over 15 orders of birds (ten Cate and Vos 1999) as well as some mammals (Kendrick et al. 1998; Montero et al. 2013) and fish (Verzijden and ten Cate 2007; Kozak and Boughman 2009; Verzijden and Rosenthal 2011). A few empirical studies have explicitly tested for a connection between sexual imprinting and sexual isolation between closely related populations or species. For example, benthic and limnetic sticklebacks sexually imprint on paternal traits under ecologically divergent selection, which results in significant sexual isolation between the two morphs (Kozak et al. 2011). Other studies in cichlids (Verzijden and ten Cate 2007), tits (Slagsvold et al. 2002), and Darwin’s finches (Grant and Grant 1997) have demonstrated that sexual imprinting can maintain sexual isolation. Therefore, sexual imprinting seems to be an important, but underexplored, avenue to speciation.

Here we assess the role of sexual imprinting in generating reproductive isolation between two mammalian species, the white-footed mouse (*Peromyscus leucopus*) and its sister species, the cotton mouse (*P. gossypinus*), which diverged in allopatry during the Pleistocene (Blair 1950). *P. leucopus* is distributed across the Midwest and eastern United States, whereas *P. gossypinus* is restricted to the Southeast (Figure 1); their ranges overlap in the Gulf Coast states, from Texas to Virginia. These species show some level of sexual isolation: when allopatric or sympatric *P. leucopus* and *P. gossypinus* are placed in large arenas, both species mate with conspecifics (Bradshaw 1965, 1968). While assortative mating in laboratory studies is potentially strong, there is mixed evidence as to whether it is strong enough to prevent hybridization in wild sympatric populations (Howell 1921; Dice 1940; McCarley 1954a; Price and Kennedy 1980; Robbins et al. 1985; Barko and Feldhamer 2002).

**Figure 1.**
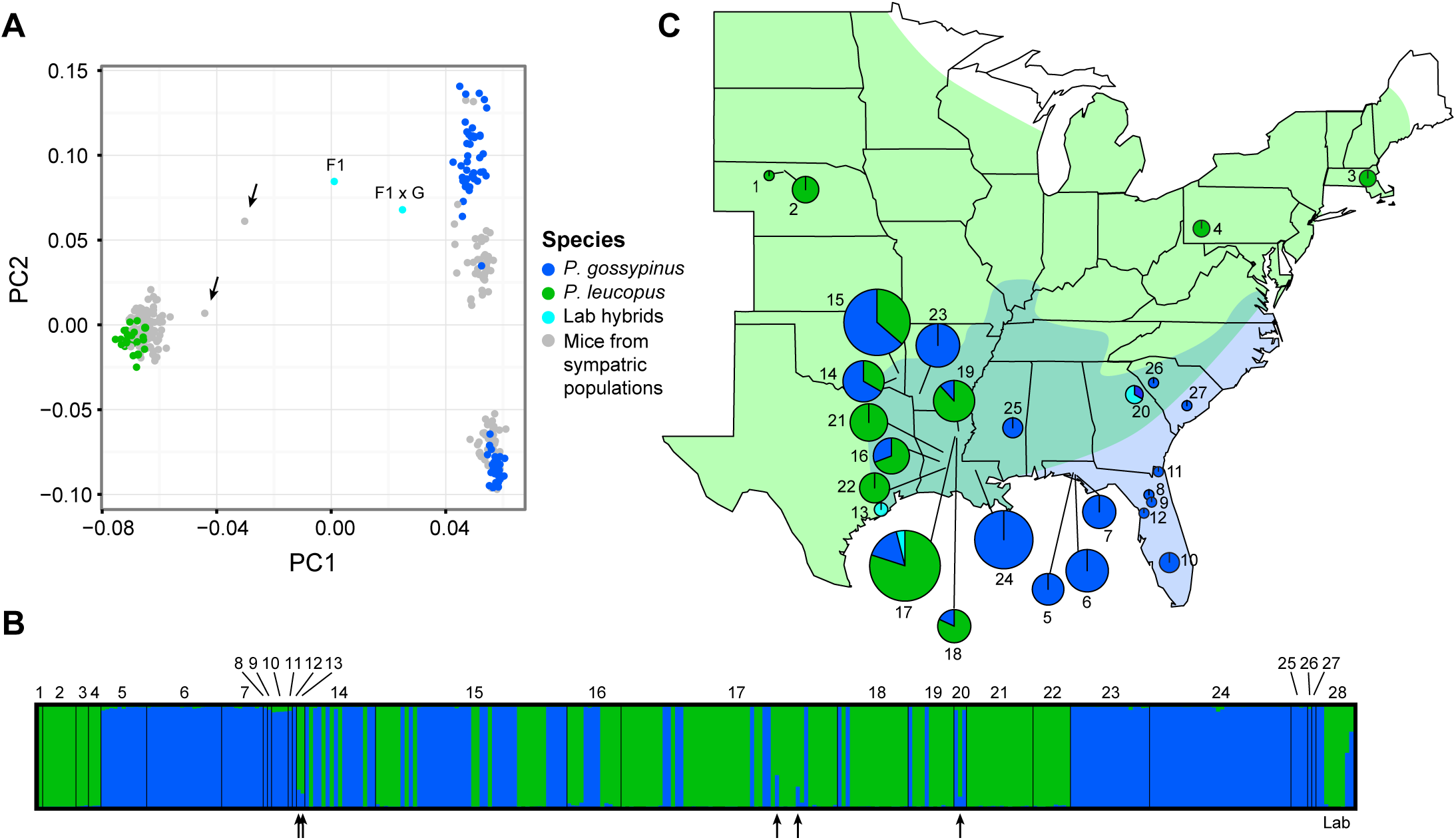
Hybridization is rare between sympatric *P. leucopus* and *P. gossypinus* mice. (**A**) Genetic PCA discriminates between species. The first PC strongly separates species based on known *P. leucopus* (green dots) and *P. gossypinus* (blue dots) mice. The second PC detects population structure within *P. gossypinus* that largely corresponds to mice collected east (higher values) and west (lower values) of the Mississippi river. Known lab-generated F1 and backcross (F1 x *P. gossypinus*) hybrids (cyan dots) fall intermediate along PC1. Mice collected from sympatry (grey dots) cluster discretely with *P. leucopus* or *P. gossypinus* with the exception of two mice that may be hybrids (arrows), but showing greater *P. leucopus* ancestry. (**B**) A Bayesian admixture model implemented in Structure also supports the partitioning of allopatric and sympatric mice into two clusters corresponding to *P. leucopus* (green) and *P. gossypinus* (blue). Individuals are represented by vertical bars showing their estimated ancestry proportions from each species. Note that Structure assigned ancestry in 28 individuals that were discarded as outliers in the genetic PCA. Populations are labeled numerically (see **C** and Supplemental Table 1 for locality information). Structure identified the same two individuals from site 17 as hybrids, but also indicated that three individuals from sites 13 and 20 may also be hybrids (arrows); however these individuals were discarded as outliers in the genetic PCA. (**C**) Range map of the two species: *P. leucopus* (green) and *P. gossypinus* (blue) adapted from (Hall and Kelson 1959; Hall 1981), showing areas of allopatry and sympatry. Pie diagrams show collecting locations and frequencies of each species scaled in size to represent the number of mice sampled at each site. For more information, see Supplemental Table 1. Mice were classified as *P. leucopus* (green), *P. gossypinus* (blue), or potential hybrids (cyan) based on the genetic PCA (shown in **A**) and Structure analysis (shown in **B**).

In this study, we used genomic data to first assess hybridization in the wild and found that the two species remain genetically distinct in sympatry despite rare hybridization events. We then measured the degree of sexual isolation between *P. leucopus* and *P. gossypinus* in the lab, and tested if it had a learned or genetic basis. Our results show that sexual imprinting produces strong sexual isolation, and suggest that learning disproportionately contributes to the total reproductive isolation we observed between two interfertile, sympatric sister species.

## METHODS

### Study species

*Peromyscus leucopus* and *P. gossypinus* are sister species that are thought to have diverged during the Pleistocene over the last 2 million years (Blair 1950; Platt et al. 2015). Fossils of *P. gossypinus* have been found in Florida and Texas (Wolfe and Linzey 1977), and *P. leucopus* fossils have been found between Texas and Pennsylvania, and as far west as Missouri (Lackey et al. 1985)—mirroring the current ranges of both species (Figure 1). The average genetic distance (*D*; Nei 1972), a proxy for divergence time, between *P. leucopus* and *P. gossypinus* is estimated to be 0.178 (Zimmerman et al. 1978). This estimate is lower than that of well-differentiated *Peromyscus* species (*D* = 0.334-0.431; Zimmerman et al. 1978), suggesting that *P. leucopus* and *P. gossypinus* are at an intermediate stage of speciation.

#### Wild samples

During April 2008 and January-February of 2010 and 2011, we collected 238 mice from ten allopatric locations and 12 sympatric locations in the central and eastern United States (Figure 1). At each location, we placed up to 300 Sherman traps every 20 feet in transects of 50 traps per line. From each mouse captured, we took liver or tail tissue and stored tissues in 100% ethanol for subsequent DNA extraction. We augmented our own sampling with tissues from museum specimens at the Harvard Museum of Comparative Zoology, Florida Museum of Natural History, Oklahoma State University Collection of Vertebrates, Sam Noble Museum Oklahoma Collection of Genomic Resources, and the Museum of Texas Tech University Genetic Resources Collection. Collecting locations and sample sizes for all animals included in this study are provided in Supplemental Table 1.

#### Lab strains

We obtained *P. leucopus* animals from the *Peromyscus* Genetic Stock Center (University of South Carolina). The *P. leucopus* stock was established with 38 founders caught between 1982-1985 from Avery County, North Carolina. In 2009, we established a stock of *P. gossypinus* animals from 18 founders caught in Jackson and Washington Counties, Florida. Both stocks were derived from allopatric sites, in which only one of the two species was present. In captivity, breeding colonies have been deliberately outbred to minimize inbreeding and preserve genetic diversity.

All animals were housed in standard mouse cages in either mated pairs (one female and one male) or in same sex cages with a maximum of five adults. Offspring were weaned into same sex cages 23 days after birth. We set the light cycle to 14 hours of light and 10 hours of dark and maintained a room temperature between 70 and 77 degrees Fahrenheit. All mice were fed a regular Purina diet (Purina Iso Pro 5P76) *ad libitum*.

In addition to maintaining these two species, we also bred hybrids in the laboratory. First generation (F1) hybrids were generated from both *P. gossypinus* female x *P. leucopus* male matings as well as the reciprocal cross. These F1 hybrids were then backcrossed to either *P. gossypinus* or *P. leucopus*.

### Detection of hybrids in sympatric populations

#### ddRADseq library construction and genotyping

We extracted genomic DNA from 374 wild-caught individuals and two lab-raised hybrids using an Autogen kit and AutoGenprep 965 instrument. We prepared double digest restriction-associated DNA tag (ddRAD) libraries from each individual following the protocol described in Peterson et al. (2012). Briefly, we digested 100-200 ng of DNA from every individual with two restriction enzymes, EcoRI-HF and MspI (New England Biolabs), and purified the reactions with AMPure XP beads (Beckman Coulter Genomics). After quantifying the cleaned and digested product on a spectrophotometer plate reader (SpectraMax Gemini XS Plate Reader), we ligated approximately 50 ng of digested DNA to uniquely barcoded EcoRI adapters and MspI adapters in a 40 µl reaction volume with T4 DNA ligase (New England Biolabs). We pooled equal amounts of 32-48 ligated samples and used two rounds of AMPure XP bead purification to reduce the total pooled volume to 30 µl. We loaded each ligation pool onto a 2% agarose Pippin Prep cassette (Sage Science) and selected fragments with a size of 300 ± 35 bp. We ran five replicate Phusion PCRs according to the Finnzymes kit directions (Thermo Fisher Scientific) for 12 cycles with 5 µl of eluted Pippin Prep product as template. Each PCR was indexed using a unique reverse primer (primer and index sequences from Peterson et al. 2012). Following PCR, we pooled all replicate reactions and purified them with AMPure XP beads to concentrate each ddRAD library. We multiplexed ddRAD libraries in equimolar ratios and sequenced 32-48 individuals per lane on the Genome Analyzer II or multiple sets of 48 individuals on the HiSeq2000 across 9 total lanes on 7 flowcells. All reads were single end and ranged between 37-47 bp.

We demultiplexed reads and aligned them by sample to a draft genome sequence of *Peromyscus maniculatus* (NCBI: GCA_000500345.1) with STAMPY run in hybrid mode using the BWA mem algorithm with default parameters (Lunter and Goodson 2011). We identified and removed adapter sequences with Picard-tools 1.100 (http://broadinstitute.github.io/picard). We realigned potential indels with the Genome Analysis Tool Kit v. 3.2-2 (GATK) IndelRealigner (McKenna et al. 2010) and performed SNP discovery across all samples simultaneously using the GATK UnifiedGenotyper (DePristo et al. 2011). We filtered alignments, keeping regions with 100 or more total reads and an average base quality greater than 20. We retained biallelic SNPs with a minimum mapping quality of 30 that were present in at least 90% of our individuals at a depth of 10 or greater. To reduce linkage among SNPs in our dataset, we identified “clusters” of SNPs within 100 bp of each other and more than 100 bp from another SNP, and we randomly selected one SNP per cluster. Our final dataset contained 3,707 SNPs and 316 mice that had over 90% of genotypes present at these SNPs (Supplemental Table 1). On average, each individual had calls for 3,607 SNPs with an average a depth of coverage of 18.6. Of these mice, we considered 71 to be of known ancestry: 20 *P. leucopus* were caught at allopatric sites or lab-raised, 49 *P. gossypinus* were caught in allopatric sites or lab-raised, and two individuals were lab-reared hybrids from our colonies. The remaining 245 individuals were of unknown ancestry and collected in the predicted sympatric range.

Short read data were deposited in GenBank (accession number: SRP123258).

#### Identification of hybrids

We first used a model-free genetic principal component analysis (PCA) to evaluate admixture between *P. leucopus* and *P. gossypinus*. We implemented genetic PCA using smartpca from the Eigensoft v.6.0.1 package (Patterson et al. 2006) and output the first ten principal components (PCs). After excluding outlier individuals and SNPs, our final dataset contained 288 individuals and 2,528 SNPs. We included individuals with known ancestry (i.e. from allopatric sites in their range or taxonomically verified museum specimens) to identify PC values corresponding to each species and identified hybrids as individuals with intermediate values along the first principal component (McVean 2009). We assessed PC significance using Tracy-Widom statistics (Patterson et al. 2006) implemented using twstats in Eigensoft v.6.0.1.

In a complementary model-based analysis, we used the Bayesian admixture model in Structure v.2.3.4 (Pritchard et al. 2000) to assign individual coefficients of membership to discrete clusters. We ran Structure with a burn-in period of 50,000 MCMC iterations, followed by 50,000 iterations, and estimated membership coefficients in five replicate runs for cluster sizes (*K*) ranging between 1 and 10. We used the Evanno method (Evanno et al. 2005) implemented in Structure Harvester (Earl and VonHoldt 2011) to determine the most likely number of clusters. We then used the full search algorithm in CLUMPP v.1.1.2 (Jakobsson and Rosenberg 2007) to estimate individual membership coefficients for all 316 individuals in our dataset across the replicate Structure runs. We considered individuals to be putative hybrids if they had >10% membership to a second cluster. To visualize our date, we used distruct v.1.1 (Rosenberg 2004).

### Measurement of sexual isolation between species

Using our laboratory *P. leucopus* and *P. gossypinus* stocks, we first tested for intrinsic postzygotic isolation and estimated sexual isolation without mate choice. We then compared our sexual isolation estimate from no-choice assays to those with mate choice to quantify the contribution of mating preferences to reproductive isolation between *P. leucopus* and *P. gossypinus*.

#### Intrinsic postzygotic isolation and sexual isolation without choice

We tested for intrinsic postzygotic sexual isolation and sexual isolation between lab-raised *P. leucopus* and *P. gossypinus* using no-choice trials. We set up 20 crosses for each conspecific and heterospecific pairing: L♀ x L♂, G♀ x G♂, L♀ x G♂, and G♀ x L♂ (in which “L” represents *P. leucopus* and “G” represents *P. gossypinus*). When F1 offspring were produced, we used these mice in additional no-choice trials in backcross mating pairs: F1♀ x L♂, F1♀ x G♂, L♀ x F1♂, and G♀ x F1♂. We avoided any sib-sib or sib-parent pairings.

We set up mating pairs by adding a sexually receptive virgin female to the cage of a virgin, sexually mature male. We determined female sexual receptivity through vaginal lavage and considered a female to be receptive between proestrus and estrus stages. We gave pairs 60 days to produce a litter, which is approximately 12 estrous cycles (mean estrous cycle length for both species is 5-6 days; Dewsbury et al. 1977) or opportunities for successful reproduction. We considered the production of offspring as a successful mating event and inferred the latency to the first successful mating by subtracting the average gestation period—23 days in both species (Pournelle 1952; Wolfe and Linzey 1977; Lackey et al. 1985)—from the total number of days until a litter was born. Although our metric for mating success is conservative because it is confounded with any fertility differences that might exist among individuals or between the species, our assay nonetheless captures hybridization between these species.

We first used the no-choice assays to test hybrid viability and fertility in our laboratory strains of *P. leucopus* and *P. gossypinus*. We scored offspring survival to reproductive age in heterospecific crosses (L♀ x G♂, G♀ x L♂), and then used these F1 hybrids in backcrosses to look for evidence of reduced fertility relative to conspecific crosses. To compare the proportion of successful mating events between conspecific and heterospecific crosses, we used a logistic regression to quantify the effects of the female species, male species, or the interaction between female and male species. We then selected the best-fit model using backward stepwise selection based on the lowest Akaike Information Criterion (AIC). We compared the 95% confidence intervals for the mean mating success among backcross pairs (F1♀ x L♂, F1♀ x G♂, L♀ x F1♂, G♀ x F1♂) to those of conspecific crosses. Together, these no-choice data provide an estimate of hybrid viability and relative fertility.

We next tested for differences in mating latency between conspecific, heterospecific, and backcross mating pairs using a non-parametric Kruskal-Wallis rank sum test followed by pairwise Wilcoxon tests with Bonferroni-corrected p-values. To quantify sexual isolation, we counted the number of successful mating events to estimate a isolation index, I_PSI_ (Rolán-Alvarez and Caballero 2000), which compares observed to expected mating events (assuming random mating among individuals) among conspecific and heterospecific pairs. This index ranges from - 1 (all mating occurred between species) to +1 (all mating occurred within species), with a value of 0 indicating equal mating among pair types. We used the number of conspecific and heterospecific pairs that produced litters to estimate I_PSI_ in JMATING v.1.0.8 (Carvajal-Rodriguez and Rolan-Alvarez 2006). We used 10,000 bootstrap replicates to estimate the sexual isolation indices, their standard deviation, and to test the hypothesis that our estimates of the sexual isolation index deviated significantly from a null hypothesis of random mating.

#### Sexual isolation with choice

We contrasted our estimate of the sexual isolation index (I_PSI_) from no-choice assays to the sexual isolation index estimated from two-way choice assays. We measured conspecific mating preferences in a two-way electronically-controlled gated mate choice apparatus that consisted of three collinear rat cages, with each pair of cages separated by two RFID antennae and gates (FBI Science Gmbh; Figure 3A). Each pair of gates was programmed to allow passage depending on the identity of the mouse. Specifically, for each trial we implanted three mice with small transponders (1.4 mm x 9 mm, ISO FDX-B, Planet ID Gmbh) in the interscapular region using a sterile hypodermic implanter and programmed the gates to allow the designated “chooser” mouse (i.e. the individual whose preference we tested) to pass freely through all cages while constraining each “stimulus” mouse to the left or right cage, respectively.

With this apparatus, we tested mate preferences of males and females of each species for conspecific and heterospecific stimuli of the opposite sex. We allowed the chooser mouse— either a sexually receptive virgin female (in proestrus or estrus as determined by vaginal lavage) or a sexually mature virgin male—to acclimate to the apparatus for one day, adding food, water, used nesting material, and a hut from each stimulus mouse’s colony housing cage to the flanking cages of the apparatus. Approximately 24 hours later, we returned the chooser mouse to the center cage if it had not already nested there, closed all gates, and added stimulus mice to the two flanking cages to allow them two to four hours to acclimate to their new environment. At lights out (4:00 pm; 14:10 hour light:dark cycle), we re-opened the gates and recorded RFID readings at all antennae as well as webcam video streams from each flanking cage for two nights (~44 hours; camera model: DLINK DCS-942L). Each chooser mouse was tested once.

At the end of each trial, we parsed a log file of RFID readings and calculated chooser preference for a stimulus as the proportion of time spent with that stimulus divided by the time spent with both stimuli. We analyzed only trials in which the chooser mouse investigated both cages during the acclimation, the chooser mouse spent at least 10 minutes investigating one stimulus during the trial, and both stimuli mice were in their cages at least 75% of the trial period (we discarded 15% of trials that did not meet these criteria).

We compared the preferences of 8-11 adults (at 9-14 weeks of age) of each species and sex for conspecific and heterospecific stimuli of the opposite sex. For female-choice trials, we tested virgin female preferences for either: (1) pairs of sexually experienced males that had successfully sired offspring with a conspecific female prior to use in the two-way choice trials (*P. leucopus*, *N* = 5 trials; *P. gossypinus*, *N* = 7 trials), or (2) pairs of virgin males as stimuli (*P. leucopus*, *N* = 6 trials; *P. gossypinus*, *N* = 4 trials). Because we did not detect a significant difference in female preference based on male stimulus sexual experience (two-sided Wilcoxon rank sum test, *P. leucopus* females: *W* = 15, *p* = 1; *P. gossypinus* females: *W* = 9, *p* = 0.41), we combined female preference data from trials with sexually experienced and virgin male stimuli. For male-choice trials, we used only virgin females as stimuli.

We estimated I_PSI_ for each sex separately in JMATING v.1.0.8 (Carvajal-Rodriguez and Rolan-Alvarez 2006) because behavior of the stimuli may not be similar across male- and female-choice trials. We estimated I_PSI_ by considering the chooser and its most preferred stimulus as a “mated” pair; when we observed no mating, we replaced zero values with a 1 to allow for bootstrapping with resampling. We used 10,000 bootstrap replicates to estimate the isolation indices and test for deviation from random mating (I_PSI_ = 0).

### Testing for sexual imprinting

To determine whether conspecific mating preferences are learned in the nest, we measured the preferences of mice from each species after they had been cross-fostered—raised from birth until weaning—by parents of the opposite species. We swapped whole litters at birth between breeding pairs of *P. leucopus* and *P. gossypinus*, reducing litters to the same number of offspring if litters differed in number of pups. All cross-fostering attempts were successful, indicating that parents readily attended to unrelated offspring. We allowed cross-fostered offspring to remain with their foster parents until weaning (23 days after birth), when we separated offspring into same sex cages; this matches the life cycle of all other mice in our study. As a control, we also cross-fostered offspring within species (i.e. swapped litters between conspecific families) to partition the effects of litter transfer and foster parent species on mating preference. Although there is mixed (or incomplete) information for whether fathers contribute parental care in *P. leucopus* and *P. gossypinus* (Hartung and Dewsbury 1979; Schug et al. 1992), we cross-fostered offspring to both parents because we maintained male-female breeding pairs in our laboratory colonies of *P. leucopus* and *P. gossypinus* and aimed to compare preferences of mice from cross-fostered and non-cross-fostered trials.

We tested the mating preferences of all cross-fostered mice in the two-way gated choice assay described above. We predicted that if young mice sexually imprint on their parents, cross-fostered mice raised with the opposite species should prefer heterospecific stimuli and exhibit a weaker preference for conspecifics compared to individuals raised by their biological parents or other unrelated conspecific parents. We evaluated the effects of chooser sex and cross-fostering treatment on preferences for *P. leucopus* in each species separately using linear modeling after applying an arcsin transformation to the proportion of time spent with *P. leucopus*. To test for the possibility that the sexes within each species might react differently to cross-fostering, we considered models with and without an interaction between chooser sex and cross-fostering treatment and selected the best-fit models using backward stepwise selection based on the lowest AIC. We compared mean estimated preferences using two-sided t-tests with Bonferroni-corrected p-values.

### Assessment of two-way choice assay

We confirmed that our two-way mate choice assay accurately predicts mating preference by measuring whether the most preferred stimulus corresponded to mating events in a subset of trials in which mating occurred. We identified trials with successful mating events by either the presence of sperm in a female reproductive tract at the end of a trial or the birth of a litter three weeks later. If a female-choice trial resulted in offspring, we determined the identity of the father by genotyping both the male stimuli and the pups at two to three microsatellite markers (loci 14, 35, and 80 from Weber et al. 2010) following the protocol described in Weber et al. 2010 (*N* = 15 trials) or screening video data for copulation events (*N* = 5 trials). We tested whether the most preferred individual (as determined by the greatest proportion of association time) predicted mating success using a linear regression. We applied an arcsin transformation to association time proportions. This analysis allowed us to determine that association time is an accurate predictor of mating, and thus reflects mating preference.

## RESULTS

### Hybridization is rare in sympatric populations

Using thousands of markers across the genome summarized in a genetic PCA, we tested for evidence of hybridization between *P. leucopus* and *P. gossypinus* in sympatric populations.

We estimated ten principal components (PCs) and removed 28 outlier individuals that exceeded six standard deviations for one of the PCs. Six of the ten PCs were significant by Tracy-Widom statistics with the following eigenvalues: (1) 37.855, (2) 4.352, (3) 3.627, (4) 3.161, (5) 3.054, and (6) 2.941. Based on clustering with known allopatric and previously identified *P. leucopus* and *P. gossypinus* specimens, PC1 clearly separates *P. leucopus* (negative values) and *P. gossypinus* (positive values) (Figure 1A). As expected, a control lab-generated F1 hybrid falls at the midpoint along PC1 and a lab backcross mouse (F1 x *P. gossypinus*) falls halfway between the F1 hybrid and the mean value of *P. gossypinus* values (Figure 1A). Of the remaining sympatric mice we collected (i.e. samples not identified as outliers), all could be easily assigned to either the *P. leucopus* or *P. gossypinus* species, with only two exceptions: two mice (EHK566 and EHK572) from Big Lake Wildlife Management Area, Louisiana had intermediate values along PC1 (Figure 1A). These admixed individuals showed greater *P. leucopus* ancestry, similar to a F1 backcross or advanced backcross to *P. leucopus*.

The second PC revealed two genetically distinct *P. gossypinus* subgroups. These likely reflect genetic differences between *P. gossypinus* subspecies, *P. gossypinus gossypinus* and *P. gossypinus megacephalus*. Specifically, higher PC2 values corresponded to mice caught east of the Mississippi river—which are more likely to be *P. g. gossypinus*—whereas lower PC2 values corresponded to mice caught west of the river—which are more likely to be *P. g. megacephalus* (Wolfe and Linzey 1977). The Mississippi river is a known biogeographic barrier for many species (Soltis et al. 2006), and our data suggest that this may also be the case for *P. gossypinus*. Only one individual from the Tunica Hills wildlife management area population in Louisiana failed to fit this pattern (Figure 1A): this individual was collected east of the Mississippi river but it clustered with individuals from the western group. We did not find any evidence to suggest a similar barrier to gene flow in *P. leucopus*, but we also did not have the equivalent population-level sampling on both sides of the river. The remaining four PCs (3, 4, 5, and 6) identified population structure within *P. leucopus* (Supplemental Figure 1).

We also estimated the optimal number of clusters in our dataset using a Bayesian admixture model in Structure. This analysis provided parallel results to our genetic PCA results: two clusters (*K* = 2) were identified in our data corresponding to *P. leucopus* and *P. gossypinus* (Figure 1B) according to the Evanno method. Unlike genetic PCA, Structure estimated cluster coefficients for all individuals in our analysis (i.e. Structure included 28 individuals that were removed as outliers in the genetic PCA). We used the average individual ancestry assignments across five replicate runs to identify potential hybrid individuals; in addition to the two potential hybrids identified in genetic PCA, three additional individuals (MCZ68799, MCZ68800, and EHK144) had ancestry proportions that were 83-90% *P. leucopus* and 10-17% *P. gossypinus*. Two of these individuals were from Nannie M. Stringfellow Wildlife Management Area, Texas (Figure 1C, site 13) and one was from Hart Creek, Georgia (Figure 1C, site 20).

### *P. leucopus* and *P. gossypinus* co-occur in mosaic sympatry

Using cluster assignments based on the genetic PCA, eight of 14 sites where the species’ ranges overlap contained both species (Figure 1B, C). The other six sites contained only a single species, highlighting the patchy distribution of both species within their broadly sympatric range from Texas and Virginia.

### No evidence for intrinsic postzygotic isolation

Previous studies suggested that there is no measurable intrinsic postzygotic isolation in laboratory crosses of *P. leucopus* and *P. gossypinus* (Dice 1937). We confirmed this result in our independent lines (i.e. different spatial and temporal origin) of these two species. We first measured reproductive success within and between species in no-choice assays. Mating success was determined largely by the female (logistic regression: *β* = 1.25, SE = 0.47, *p* = 0.008; Supplemental Table 2), with *P. leucopus* females showing greater mean mating success than *P. gossypinus* (Supplemental Figure 2). Importantly, this means that *P. leucopus* females had greater reproductive success with *P. gossypinus* males (12/20 pairs had offspring) than the reciprocal cross between *P. gossypinus* females and *P. leucopus* males (6/20 pairs had offspring), indicating some asymmetry in mate preferences, copulation attempts, or female fertility.

Successful heterospecific crosses confirmed the ability to produce viable F1 hybrids, which survive until reproductive age. In addition, we compared the mating successes of backcrosses to conspecific and heterospecific mates. We found that F1 hybrids are as fertile in backcrosses (i.e. had similar frequency of litter production) as either conspecific or heterospecific crosses, and that all backcross offspring are also viable (Supplemental Figure 2).

### Mate choice leads to sexual isolation

We next examined whether mating preferences lead to sexual isolation between the species in a laboratory environment. In no-choice assays, heterospecific pairs hybridized and produced viable offspring (Supplemental Table 3), indicating no measurable sexual isolation in the absence of mate choice (I_PSI_ = 0.00, *SD* = 0.19, *p* = 0.960). However, conspecific, heterospecific, and backcross mating pairs had significantly different latencies to produce offspring (Figure 2; Kruskal-Wallis: *χ*^2^ = 6.7626, df = 2, *p* = 0.034). Pairwise comparisons between mating pairs revealed significance differences in latency to mating only between conspecific and heterospecific mating pairs (W = 69, *p*_Bonferroni_ = 0.010), but not between conspecific and backcross mating pairs (W = 130, *p*_Bonferroni_ = 0.949) or between heterospecific and backcross mating pairs (W = 188.5, *p*_Bonferroni_ = 1). Heterospecific pairs took an average of 5.4 days longer to produce litters than conspecific pairs, indicative of either delayed heterospecific mating or longer hybrid gestation times. This delay is roughly equivalent to one estrus cycle in *Peromyscus* (Dewsbury et al. 1977). No significant differences were detected between the two conspecific pair types, L♀ x L♂ or G♀ x G♂ (W = 53, *p*_Bonferroni_ = 0.238), or between the two heterospecific pair types, L♀ x G♂, and G♀ x L♂ (W = 25, *p*_Bonferroni_ = 0.645).

**Figure 2.**
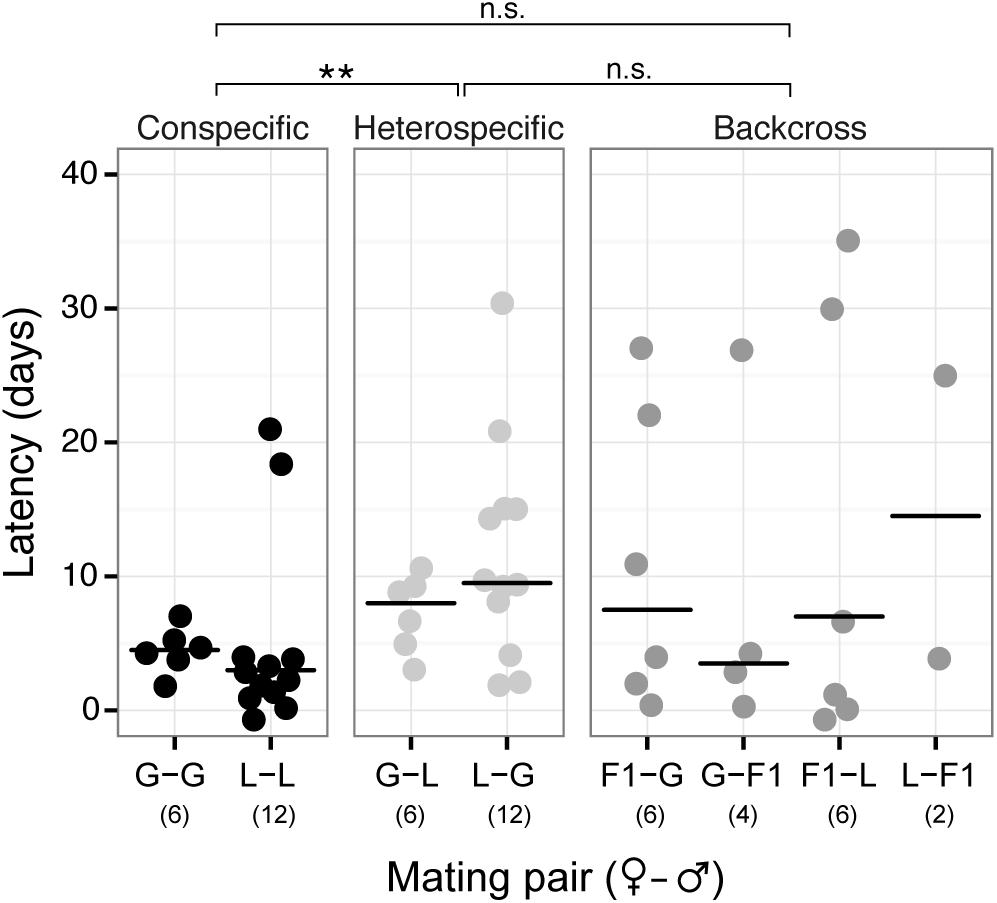
Latency to mating between *P. leucopus* (L), *P. gossypinus* (G) and their hybrids (F1). Estimated days since copulation are shown for conspecific, heterospecific, and backcross mating pairs that produced offspring (sample size in parentheses) in no-choice assays. F1 hybrids were generated with both LxG and GxL crosses. In all pairs, the female is listed first. ** *p* = 0.01.

By contrast, we detected significant sexual isolation between the species in two-way choice assays (Supplemental Table 3). Sexual isolation estimates were similar in female- and male-choice trials: *P. leucopus* and *P. gossypinus* females strongly preferred conspecific mates (Figure 3B; I_PSI_ = 0.75, *SD* = 0.14, *p* < 0.01) as did *P. leucopus* and *P. gossypinus* males (Figure 3B; I_PSI_ = 0.75, *SD*= 0.15, *p* < 0.01). More generally, there were strong preferences for conspecific mates in both species, regardless of sex.

**Figure 3.**
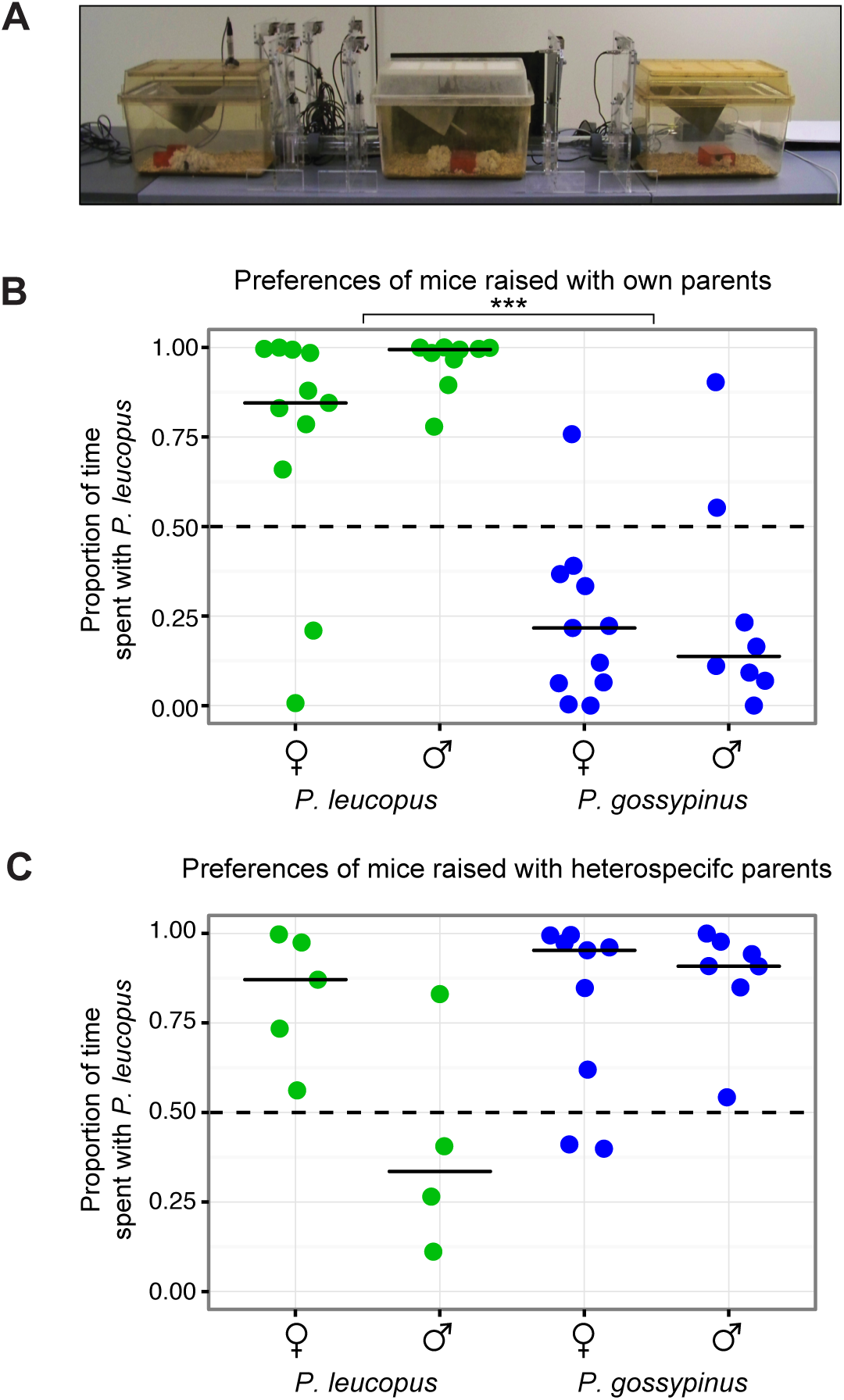
Mating preferences in two-way choice trials. (**A**) Photograph of the mate-choice apparatus. Center chamber is connected to two test chambers, each housing a “stimulus” animal, separated by gated doors activated by only the “chooser” animal. (**B**) Mating preferences for mice raised by their own parents. *P. leucopus* spent greater time with *P. leucopus* stimuli than both *P. gossypinus* sexes. (**C**) Mating preferences for mice raised by heterospecific foster parents. *P. leucopus* males were significantly affected by cross-fostering (*p* = 0.004), whereas *P. leucopus* females were not. Both *P. gossypinus* sexes spent significantly more time with the heterospecific stimulus than when raised by their own parents (*p* < 0.001).

### Sexual imprinting contributes to sexual isolation in at least one species

We then investigated whether mating preferences in these species had a learned or genetic basis using a series of cross-fostering experiments. We found that cross-fostering had different effects on mating preference in the two focal species. In *P. leucopus*, mating preference was best predicted by a full model with cross-fostering, sex, and their interaction (*F* = 5.09 on 3 and 25 df, *p* = 0.007); a reduced model was not selected by AIC (Supplemental Table 4). When raised with their own parents, *P. leucopus* of both sexes preferred *P. leucopus* stimuli (Figure 3B; estimated proportion of female time spent with *P. leucopus* = 0.689; estimated proportion of male time spent with *P. leucopus* = 0.959). *P. leucopus* males that were cross-fostered significantly changed their preference (Figure 3C; estimated proportion of cross-fostered male time spent with *P. leucopus* = 0.184; t = -3.853, *p*_Bonferroni_ = 0.003), whereas cross-fostering did not significantly change female preference (Figure 3C; estimated proportion of cross-fostered female time spent with *P. leucopus* = 0.764; t = 0.390, *p*_Bonferroni_ = 1). Thus, *P. leucopus* females always preferred *P. leucopus* to *P. gossypinus* mates, whereas a male spent more time with the species with which it was raised.

In *P. gossypinus*, mating preference was best predicted by a reduced model (Supplemental Table 5) with a significant cross-fostering term but no significant sex effects or interactions between cross-fostering and sex (*F* = 51.31 on 1 and 33 df, *p* < 0.001). When raised with their own parents, *P. gossypinus* of both sexes preferred *P. gossypinus* stimuli (Figure 3B; estimated preference for *P. leucopus* = 0.069), whereas *P. gossypinus* raised with *P. leucopus* preferred *P. leucopus* stimuli (Figure 3C; estimated preference for *P. leucopus* = 0.781).

To confirm that cross-fostering affect was caused by the foster parent species and not due to transferring litters, we collected an additional control dataset for *P. gossypinus.* We cross-fostered *P. gossypinus* to unrelated *P. gossypinus* foster parents (females: *N* = 4, males: *N* = 7) and found that foster species, and not the transfer itself, affected *P. gossypinus* preferences (Supplemental Figure 3). Pairwise t-tests on arcsin-transformed proportion of time spent with *P. leucopus* revealed no significant differences between *P. gossypinus* raised with their own parents or unrelated conspecific parents (*t* = -0.72, df = 15.38, *p*_Bonferroni_ = 1).

To examine the effects of sexual imprinting on sexual isolation, we calculated the sexual isolation index (I_PSI_) assuming the most preferred stimulus from each heterospecific cross-fostered trial (Figure 3C) as a “successful mating”. Cross-fostering eliminated sexual isolation in female-choice trials (I_PSI_ = 0.25, *SD* = 0.34, *p* = 0.57) and male-choice trials (I_PSI_ = -0.29, *SD* = 0.42, *p* = 0.32). Thus, our cross-fostering results confirm that sexual isolation between *P. leucopus* and *P. gossypinus* is the result of sexual imprinting.

### Two-way choice test accurately measures preference

To confirm that the time spent with a stimulus mouse was an accurate predictor of mate preference and hence mate choice, we recorded 20 mating events in our two-way choice assays: 12 mating events occurred in trials where choosers were raised with their own parents and 8 occurred in trials where choosers were raised with heterospecific foster parents. In 19 out of 20 trials, choosers mated with the stimulus individual with whom they spent the most time (Figure 4). Mating outcome (with conspecific or heterospecific stimulus) was predicted by the proportion of time spent with the conspecific stimulus (logistic regression: *β* = 10.06, *SE* = 4.86, *p* = 0.04), indicating that our two-way choice assay accurately detects mating preferences.

**Figure 4.**
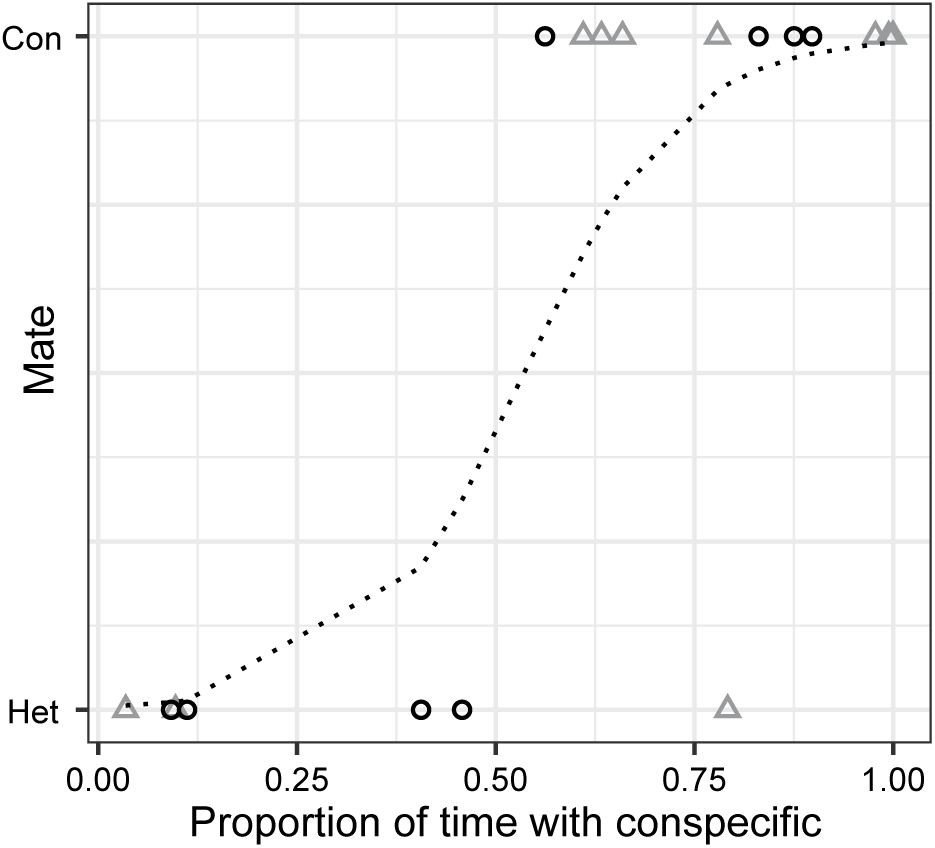
The proportion of time spent with a stimulus predicts mating outcome in trials when mating occurred. Mating occurred in 12 trials when choosers were raised with their own parents (gray triangles) and 8 trials in where choosers were raised with heterospecific parents (black dots). Dotted line indicates the predicted probability for mate choice (conspecific versus heterospecific) given the proportion of time a chooser spent with a conspecific individual, which strongly predicts the mating partner (*p* = 0.038). With the exception of one *P. leucopus* female raised with her own parents, all mice spent more time with their preferred mate.

## DISCUSSION

Sexual imprinting can be a powerful generator of sexual isolation because it quickly and effectively associates preferences with traits in populations. Furthermore, sexual imprinting has been documented in a diversity of taxa—e.g. birds, fish, mammals, amphibians, and insects— suggesting it may be a broadly important driver of speciation (Immelmann 1975). Our study shows that that sexually imprinted mate-choice has likely contributed to and maintained strong sexual reproductive isolation between a pair of mammalian sister species.

### Rare hybridization in sympatry indicates a high degree of reproductive isolation

To test the strength of reproductive isolation between *P. leucopus* and *P. gossypinus* in nature, we first collected mice from across their ranges and used genomic data to test for hybridization between these species in sympatry. Classic studies by mammalogists in the mid 1900’s reported conflicting results as to the extent of interspecific hybridization in sympatric populations. In Louisiana, Alabama, and southern Illinois, Howell (1921), McCarley (1954a), and later Barko and Feldhamer (2002) identified a few intermediate individuals resembling hybrids based on morphology and allozyme genotypes. By contrast, Dice (1940) found no evidence of morphological intermediates in his studies in Virginia. Thus, the degree of hybridization if any between these two species in the wild has been contested historically.

In total, our analyses identified only five potential wild hybrids out of 245 mice that were collected from locales where the species’ ranges overlap (Figure 1C). Two hybrids were identified in both genetic analyses (genetic PCA and Structure) and three identified by Structure alone; all had greater proportions of *P. leucopus* ancestry. Thus, we found that approximately 2% of individuals were admixed. Interestingly, the five hybrids we identified occurred in locations where *P. gossypinus* was the rarer species, providing one explanation as to why they likely backcrossed to *P. leucopus*. Nonetheless, both model-free and model-based clustering methods showed that the vast majority of mice in our study clustered into two discrete groups, one for each species, regardless of population. Our genomic analysis thus suggests that, despite rare hybrids, *P. leucopus* and *P. gossypinus* remain genetically distinct in nature.

Our genomic data, which allowed us to confidently assign individuals to species, also revealed that *P. leucopus* and *P. gossypinus* are distributed in a mosaic sympatry, with several sites containing only one species (six of 14 sampling sites). This patchiness could be driven by differences in microhabitat use: *P. leucopus* often occupy upland habitat and use more arboreal nest sites while *P. gossypinus* often occupy swamps and bottomland habitat and use more ground nest sites when they co-occur (McCarley 1954b, 1963; Taylor and McCarley 1963). However, these habitat differences are not enough to exclude contact in sympatry because both species can be trapped in the same patch of forest, especially where these habitat types abut (Dice 1940; Calhoun 1941; Price and Kennedy 1980; Roehrs et al. 2012). In fact, we often caught both species in the same trap line, indicating that the species overlap within each other’s cruising ranges. Similarly, there do not appear to be any significant differences in breeding seasons: the two species have overlapping peak reproductive activities in the winter months, but adults from both species can also be caught in reproductive condition throughout the year in Texas, Louisiana, and Alabama (Pournelle 1952; McCarley 1954c; Wolfe and Linzey 1977). Thus, the distributions, habitat preferences, and breeding seasons are unlikely to form complete or even strong reproductive barriers, suggesting that behavioral differences may be an important contributor to the level of reproductive isolation we observed in the wild.

### Learned sexual isolation in *P. leucopus* and *P. gossypinus*

As previous studies suggested that mating preferences might explain the lack of hybridization in the wild, we tested for evidence of sexual isolation. Using no-choice and choice assays to examine *P. leucopus* and *P. gossypinus* mating preferences, we found that conspecific preferences form a significant sexual barrier between the two species. Without a choice of mates, *P. leucopus* and *P. gossypinus* did not show significant sexual isolation, although there was an increase in latency to mate in heterospecific crosses relative to conspecific crosses. However, when given a choice of mates, the species mated assortatively, and we estimated the average sexual isolation index (I_PSI_) between the species to be 0.651. While sexual isolation is high, it is not yet complete (I_PSI_ < 1) between these species. However, the amount of sexual isolation we have observed is far greater than what has been detected among cactophilic (I_PSI_ = 0.12; Etges and Tripodi 2008) or Caribbean *Drosophila* (I_PSI_ = 0.159-0.282; Yukilevich and True 2008), walking stick insect populations (I_PSI_ = 0.24-0.53; Nosil et al. 2013), or gold and normal Nicaraguan cichlid color morphs (I_PSI_ = 0.39 and 0.86; Elmer et al. 2009), placing *P. leucopus* and *P. gossypinus* quite far along a speciation continuum.

Using cross-fostering experiments, we found that conspecific mating preferences were largely determined by sexual imprinting. This result implies that sexual isolation, a primary reproductive barrier between sympatric, interfertile populations of *P. leucopus* and *P. gossypinus*, is mostly due to learning. This work also implies that there are informative cues that the species reliably use to distinguish between *P. leucopus* from *P. gossypinus* (but we do not yet know if these signals are chemical, audial, or visual). Our work suggests that mammalian species that sexually imprint might therefore be poised to form strong reproductive barriers at earlier stages in the speciation process that enable sympatry without rampant hybridization. In fact, other species of *Peromyscus* are also affected by cross-fostering (Carter and Brand 1986; Bester-Meredith and Marler 2001), raising the possibility that their speciation trajectories could have similarly been affected by learned mating preferences.

Intriguingly, our cross-fostering studies also revealed that the degree of imprinting differed by species and sex. We found that both male and female *P. gossypinus* strongly sexually imprinted on their foster parent species. By contrast, we found that *P. leucopus* also sexually imprint on parents, although only weakly. Some *P. leucopus* males had a reduced preference for conspecifics when raised with heterospecific parents, whereas all *P. leucopus* females appeared unaffected by cross-fostering. *P. leucopus* showed a similar sexual difference in a study that examined preferences for soiled bedding after cross-fostering to grasshopper mice, *Onychomys torridus* (McCarty and Southwick 1977): although both male and female *P. leucopus* raised with *O. torridus* parents had decreased preference for conspecific soiled bedding, the effect was more dramatic in males than females. Thus, both *P. leucopus* and *P. gossypinus* appear to learn mating preferences, but the degree of sexual imprinting varies between the two species, and between the sexes in *P. leucopus*.

### Interspecific and sex-biased differences in sexual imprinting

While *P. gossypinus* males and females form strong conspecific mating preferences through sexual imprinting, only males of its sister species, *P. leucopus*, appear to sexually imprint. Such asymmetric effects of sexual imprinting on congeneric species may not be usual. For example learning affects mating preferences asymmetrically in congeneric tits (Slagsvold et al. 2002) and swordtails (Verzijden et al. 2012b). What might cause this variation in learning between *Peromyscus leucopus* and *P. gossypinus,* and why are preferences in *P. leucopus* females robust to sexual imprinting?

One possibility is that conspecific mating preferences are innate and genetically controlled in *P. leucopus* females due to reinforcement with *Peromyscus maniculatus*, a sympatric species whose geographic range largely overlaps with *P. leucopus* (Hall 1981). Because hybrids between *P. leucopus* and *P. manciulatus* are inviable (Maddock and Dawson 1974), natural selection could have reinforced the canalization of conspecific mating preferences in *P. leucopus* females if they incur high costs from heterospecific mating. Innate genetic conspecific mating preferences in *P. leucopus* females would suggest that the hybrids we detected are more likely to been progeny from crosses between *P. leucopus* males with *P. gossypinus* females.

Alternatively, *P. leucopus* may sexually imprint on parents but modify their preferences after interactions with conspecifics and heterospecifics. In our study, male *P. leucopus* stimuli may direct more copulatory behavior toward *P. leucopus* females, whereas male *P. gossypinus* stimuli may be more antagonistic, thereby causing females to reverse learned preferences for heterospecifics. Such preference reversals following cross-fostering have been observed in other species (Rosenthal 2017). For example, a study of the effects of cross-fostering between sheep and goats found that females raised with heterospecific foster parents initially preferred heterospecific males, but later preferred conspecifics after a year of socialization (Kendrick et al. 1998); in contrast, males continued to prefer mates of their foster parent species. Similarly, female zebra finches cross-fostered with Bengalese foster parents spent more time with Bengalese males but directed more sexually receptive tail quivering behavior to conspecific males who sang more vigorously and frequently (ten Cate and Mug 1984). If mating preferences in *P. leucopus* females are indeed learned but susceptible to adult social interactions, mating attempts by *P. gossypinus* males might account for the few hybrids we observed in our study.

Finally, the species and sexes could differ in their sexual imprinting sets. Imprinting on fathers is more likely to evolve than imprinting on mothers (Tramm and Servedio 2008) and could potentially occur in *Peromyscus,* as it does in *Mus* (Montero et al. 2013), if males associate with juvenile offspring. Should the few hybrids we discovered be primarily produced from one type of heterospecific cross, imprinting on either mothers or fathers would lead to biased introgression. In addition, imprinting on siblings is also possible given that we cross-fostered whole litters to male-female pairs. Thus, the own-species bias in *P. leucopus* females but not *P. gossypinus* might also be the result of imprinting on siblings. Future experiments could experimentally test for the imprinting set, and even specific cues involved, determining if and how they differ between species and sexes.

### Reproductive isolation in sympatry

Sexual imprinting could be even stronger between *P. leucopus* and *P. gossypinus* than what we have measured in the lab if it were reinforced in sympatric populations (Irwin and Price 1999; Servedio et al. 2009). Although we did not find evidence of hybrid inviability or sterility in the laboratory using allopatric stocks, the degree of hybrid fertility could vary in severity in natural hybrid zones (e.g. Turner et al. 2011). Additionally, extrinsic postzygotic barriers, such as behavioral sterility, may create an opportunity for reinforcement. Previous work found that *P. leucopus* and *P. gossypinus* reciprocal hybrids initiated copulation less frequently than either *P. leucopus* or *P. gossypinus* despite having similar copulatory behaviors (Lovecky et al. 1979).

Hybrids also differed in exploratory behavior compared to either parental species (Wilson et al. 1976), which may reduce hybrid fitness. Finally, hybrids might be behaviorally sterile if they have intermediate mating traits. For example, hybrids between *M. m. musuculus* and *M. m. domesticus* have intermediate urinary signals that are selected against by each subspecies (Latour et al. 2014). That we have found moderate sexual isolation in our allopatric lab stocks implies that learning could be selected and strengthened in sympatry if it reduced the production of behaviorally unfit hybrids. The potential for behaviorally-induced reinforcement, coupled with the fact that moderate sexual imprinting induces sexual isolation in our lab stocks, could boost reproductive isolation in sympatry and help explain the paucity of hybrids we have observed in our study.

## CONCLUSION

Our study supports an emerging view that sexual imprinting may be vital to the generation and maintenance of sexual reproductive barriers. Pending divergence in an imprintable trait, a species that learns mating preferences may develop significant sexual isolation that might mitigate the homogenizing effects of hybridization. Our demonstration of sexual imprinting in *Peromyscus gossypinus* and *P. leucopus*, sympatric sister species that have few other measurable reproductive barriers between them, suggests that sexual imprinting may be an important contributor to their overall reproductive isolation. However, it is notable that the strength of imprinting differs between the species, and in one species, is largely sex-specific. Nonetheless, sexual imprinting could sculpt reproductive isolation in subspecies (e.g. benthic and limnetic sticklebacks) undergoing initial morphological and behavioral divergence, or help preserve reproductive isolation between already divergent species, as we see in *P. leucopus* and *P. gossypinus*. Examining the role of sexual imprinting in similar cases of speciation driven by sexual reproductive barriers will continue to expand our understanding of the role of behavior in speciation.

## Acknowledgements

We thank members of the Hoekstra lab who helped with field work: A. Chiu, G. Gonçalves, V. Domingues, E. zu Ermgassen, H. Fisher, E. Jacobs-Palmer, E. Kingsley, K. Lin, C. Linnen, H. Metz, B. Peterson, and J. Weber; we also thank E. Pivorun, R. Sikes, and their students for assistance. J. Chupasko and M. Omura from the Harvard Museum of Comparative Zoology Mammal Department helped with specimen curation. Nokuse Plantation granted us permission to trap on their property. The following museums donated tissue samples: Harvard Museum of Comparative Zoology, Florida Museum of Natural History, Oklahoma State University Collection of Vertebrates, Oklahoma Collection of Genomic Resources, and the Museum of Texas Tech University. We also wish to thank N. Delaney, K. Ferris, J. Mallet and our reviewers for helpful discussions and comments on this manuscript. This work was funded by: NSF GRFP, NSF DDIG, and grants from the American Society of Mammalogists, Animal Behavior Society, and the Harvard University Mind, Brain & Behavior Initiative to EKD. HEH is an Investigator of the Howard Hughes Medical Institute. The authors have no conflicts of interest.

---

**Supplemental Figure 1.**
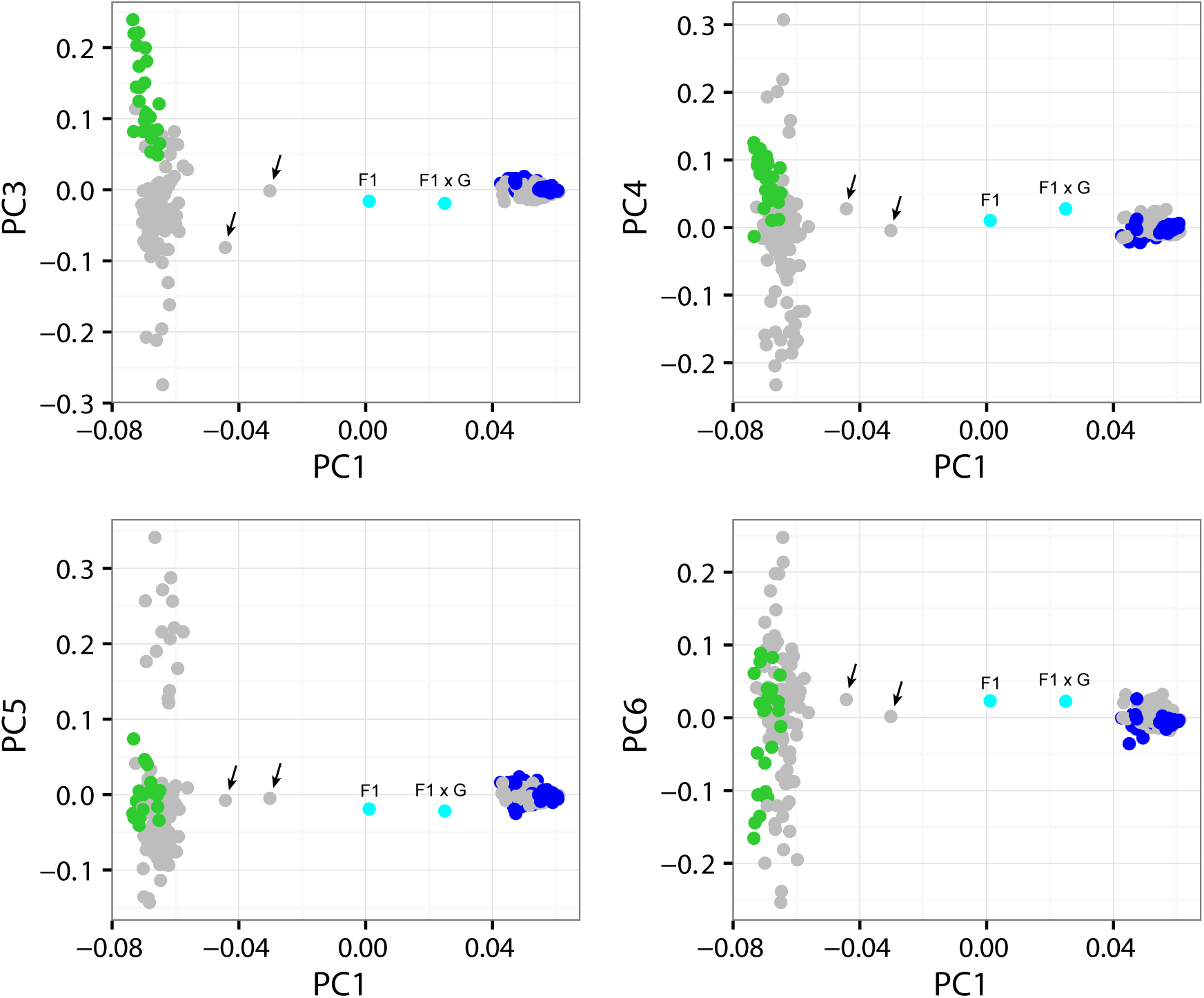
The first principal component (PC) separates the species *P. leucopus*, *P. gossypinus* and their possible hybrids. Mice collected from sympatry (gray dots) cluster discretely with either known allopatric *P. leucopus* (green dots) or known *P. gossypinus* (blue dots) with the exception of two individuals that may be hybrids (arrows) but show more *P. leucopus* ancestry. The possible hybrids fall intermediate along PC1, similar to known lab-generated F1 and backcross (F1 x *P. gossypinus*) hybrids (cyan dots). PCs 3, 4, 5, and 6 identified population structure within *P. leucopus*.

**Supplemental Figure 2.**
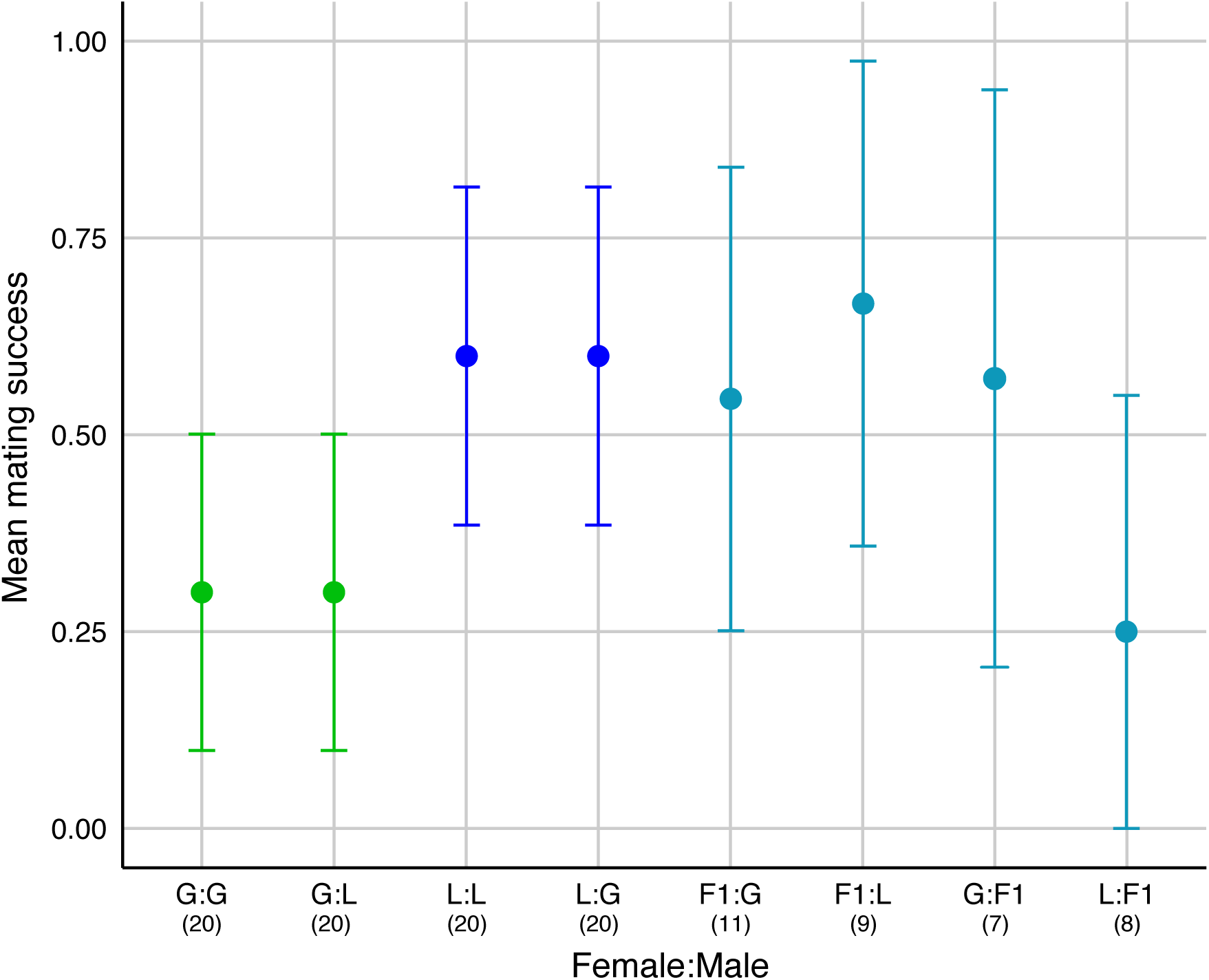
Mean proportion of mating successes, defined as the production of a litter, in no-choice trials with *P. leucopus*, *P. gossypinus*, and F1 hybrids (from L:G and G:L crosses). Proportions (dots) and their 95% confidence intervals are plotted for each cross (sample size in parentheses). Confidence intervals overlap for all cross types, indicating that hybrids do not suffer reduced mating success in backcrosses compared to conspecific crosses (G:G or L:L).

**Supplemental Figure 3.**
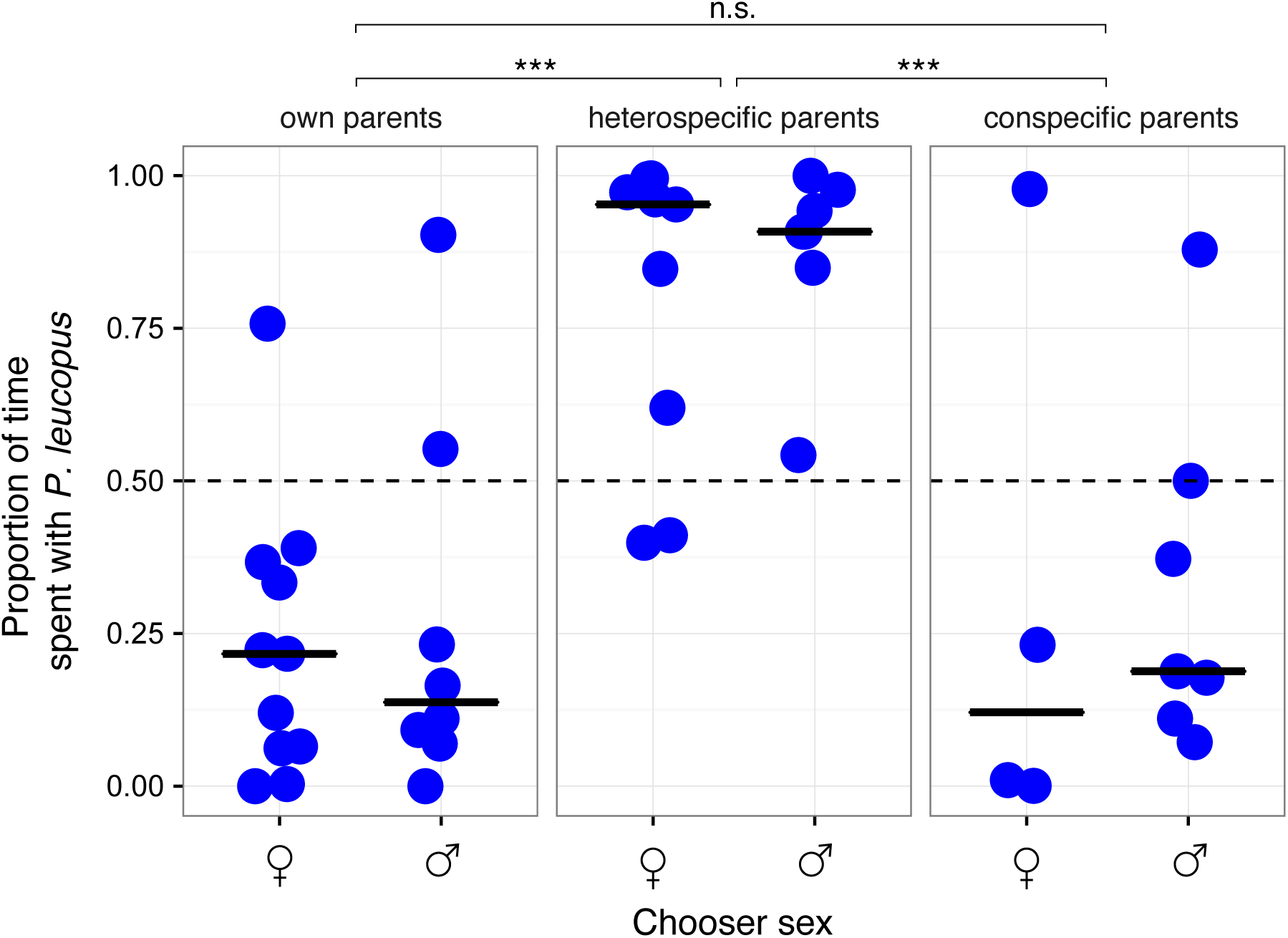
*Peromyscus gossypinus* mating preferences for mice raised with their own parents, heterospecific parents, or unrelated conspecific parents. *P. gossypinus* raised with heterospecific parents differed signifcantly from mice raised with their own parents (t = -7.04, df = 28.89, *p*_*Bonferroni*_ < 0.01) and mice raised with conspecific parents (t = 4.31, df = 18.90, *p*_*Bonferroni*_ < 0.01), but *P. gossypinus* raised with their own parents did not significantly differ from mice raised with unrelated conspecific parents (t = -0.72, df = 15.38, *p*_*Bonferroni*_ = 1).

**Supplemental Table 1.**
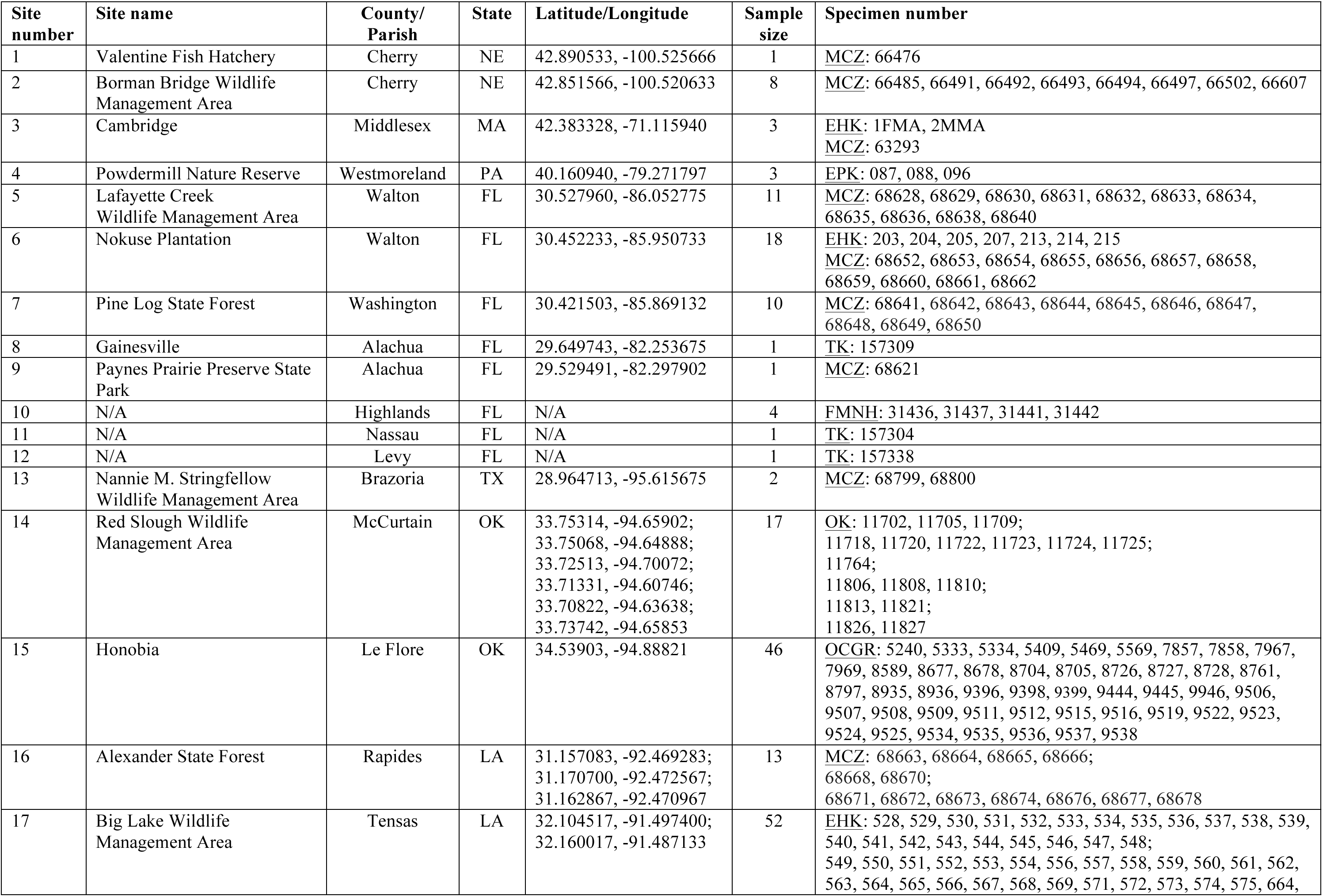
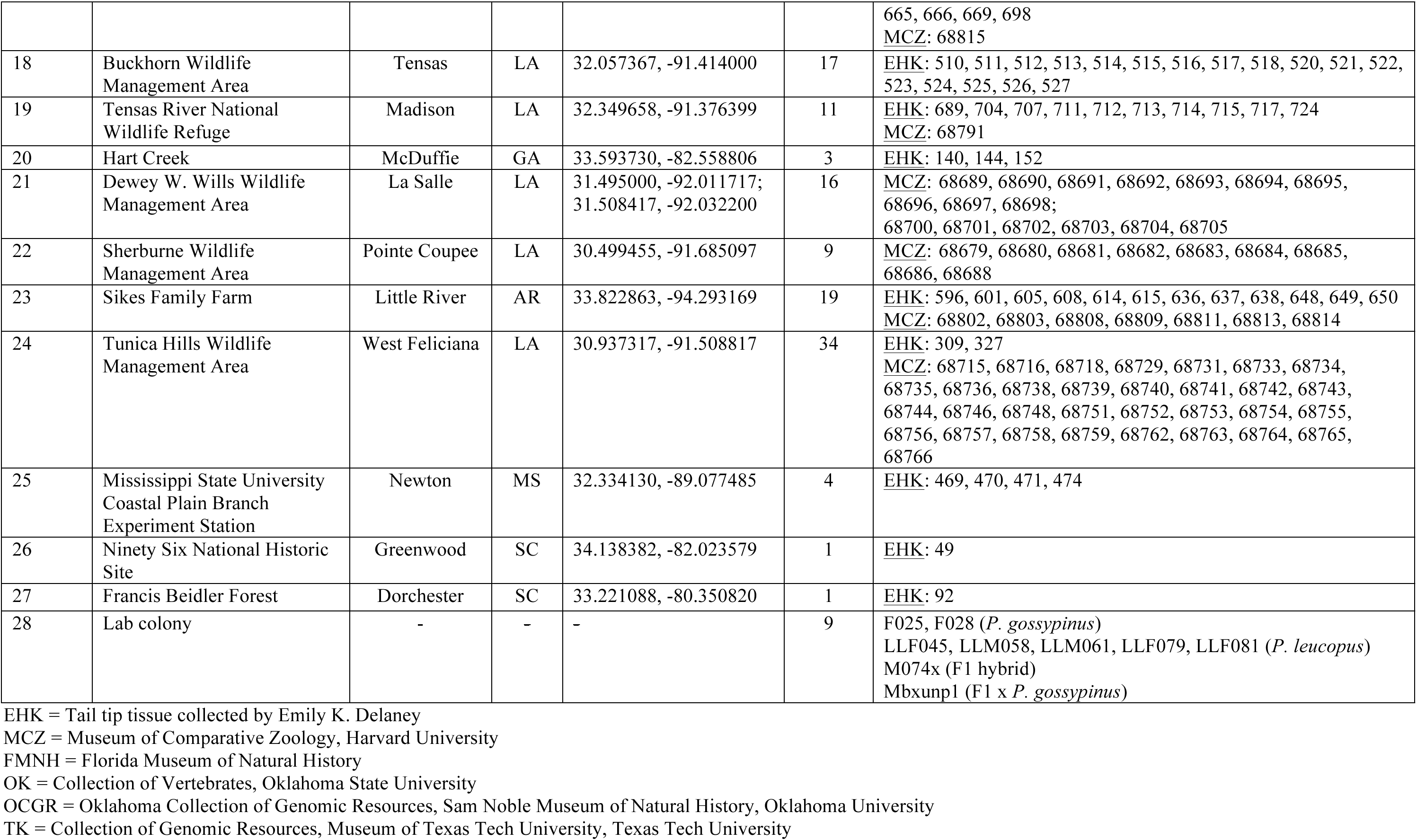
Trapping locations and specimen information.

**Supplemental Table 2.** Linear models for mating success in conspecific and heterospecific no-choice trials.

| **Model** | **Log-likelihood** | **Number of Parameters** | **AIC** | **ΔAIC** |
| --- | --- | --- | --- | --- |
| Mating success* ~<br>Female species +<br>Male species +<br>Female<br>species×Male<br>species | -51.355 | 4 | 110.710 | - |
| Mating success ~<br>Female species +<br>Male species | -51.355 | 3 | 108.710 | 2.000 |
| Mating success ~<br>Female species | -51.355 | 2 | 106.710 | 2.000 |

\*Mating success is defined as a confirmed copulation event (either the birth of a litter or presence of sperm).

**Supplemental Table 3.** Mating results for no-choice, female-choice, and male-choice trials with non-cross-fostered and cross-fostered mice.

| Trial type (Female x Male) | Matings | Number of trials |
| --- | --- | --- |
| No-choice |  |  |
| G x G | 6 | 20 |
| G x L | 6 | 20 |
| L x G | 12 | 20 |
| L x L | 12 | 20 |
| Female-choice* |  |  |
| G x G | 10 | 11 |
| G x L | 1 | 11 |
| L x G | 2 | 11 |
| L x L | 9 | 11 |
| Male-choice* |  |  |
| G x G | 6 | 8 |
| G x L | 2 | 8 |
| L x G | 0 | 9 |
| L x L | 9 | 9 |
| Female-choice (cross-fostered)* |  |  |
| G x G | 2 | 9 |
| G x L | 7 | 9 |
| L x G | 0 | 5 |
| L x L | 5 | 5 |
| Male-choice (cross-fostered)* |  |  |
| G x G | 0 | 7 |
| G x L | 7 | 7 |
| L x G | 1 | 4 |
| L x L | 3 | 4 |

\*Matings are inferred from female- and male-choice trials based on proportion of time spent with each stimulus (see Figures 3 and 4). The stimulus individual with whom the chooser spent more time was considered a mate.

**Supplemental Table 4.** Linear models for *Peromyscus leucopus* preference.

| **Model** | **Log-likelihood** | **Number of Parameters** | **AIC** | **ΔAIC** |
| --- | --- | --- | --- | --- |
| Preference* ~ Sex + CrossFoster + Sex×CrossFoster | -12.310 | 5 | 34.620 | - |

\*Preference is defined as the arcsin-transformed proportion of time spent with *P. leucopus*.

**Supplemental Table 5.** Linear models for *Peromyscus gossypinus* preference.

| **Model** | **Log-likelihood** | **Number of Parameters** | **AIC** | **Delta AIC** |
| --- | --- | --- | --- | --- |
| Preference* ~ Sex + CrossFoster + Sex×CrossFoster | -10.238 | 5 | 30.477 | 0 |
| Preference ~ ChooserSex + CrossFoster | -10.261 | 4 | 28.523 | 1.954 |
| Preference ~ CrossFoster | -10.502 | 3 | 27.004 | 1.519 |

\*Preference is defined as the arcsin-transformed proportion of time spent with *P. leucopus*.

## References

Barko, V. A., and G. A. Feldhamer. 2002. Cotton mice (*Peromyscus gossypinus*) in southern Illinois: evidence for hybridization with white-footed mice (*Peromyscus leucopus*). Am. Midl. Nat. 147:109–115.

Bester-Meredith, J. K., and C. A. Marler. 2001. Vasopressin and aggression in cross-fostered California mice (*Peromyscus californicus*) and white-footed mice (*Peromyscus leucopus*). Horm. Behav. 40:51–64.

Blair, W. F. 1950. Ecological factors in speciation of *Peromyscus*. Evolution 4:253–275.

Boughman, J. W., H. D. Rundle, and D. Schluter. 2005. Parallel evolution of sexual isolation in sticklebacks. Evolution 59:361–373.

Bradshaw, W. N. 1965. Species discrimination in the *Peromyscus leucopus* group of mice. Texas J. Sci. 17:278–293.

Bradshaw, W. N. 1968. Progeny from experimental mating tests with mice of the *Peromyscus leucopus* group. J. Mammal. 49:475–480.

Calhoun, J. B. 1941. Distribution and food habits of mammals in the vicinity of the Reelfoot Lake Biological Station. J. Tennessee Acad. Sci. 16:177–187.

Carter, R. L., and L. R. Brand. 1986. Species recognition in wild-caught, laboratory-reared and cross-fostered *Peromyscus californicus* and *Peromyscus eremicus* (Rodentia, Cricetidae). Anim. Behav. 34:998–1006.

Carvajal-Rodriguez A., and E. Rolan-Alvarez. 2006. JMATING: a software for the analysis of sexual selection and sexual isolation effects from mating frequency data. BMC Evol. Biol. 6:40.

Coyne, J. A., and H. A. Orr. 1989. Patterns of speciation in *Drosophila*. Evolution 43:362–381.

Coyne, J. A., and H. A. Orr. 1997. “Patterns of speciation in *Drosophila*” revisited. Evolution 51:295–303.

DePristo, M. A., E. Banks, R. Poplin, K. V Garimella, J. R. Maguire, C. Hartl, A. A. Philippakis, G. del Angel, M. A. Rivas, M. Hanna, A. McKenna, T. J. Fennell, A. M. Kernytsky, A. Y. Sivachenko, K. Cibulskis, S. B. Gabriel, D. Altshuler, and M. J. Daly. 2011. A framework for variation discovery and genotyping using next-generation DNA sequencing data. Nat. Genet. 43:491–498.

Dewsbury, D. A., D. Q. Estep, and D. L. Lanier. 1977. Estrous cycles of nine species of muroid rodents. J. Mammal. 58:89–92.

Dice, L. R. 1937. Fertility relations in the *Peromyscus leucopus* group of mice. Contrib. Lab. Vert. Genet. 4:1–3.

Dice, L. R. 1940. Relationships between the wood-mouse and the cotton-mouse in eastern Virginia. J. Mammal. 21:14–23.

Earl, D. A., and B. M. VonHoldt. 2011. STRUCTURE HARVESTER: a website and program for visualizing STRUCTURE output and implementing the Evanno method. Conserv. Genet. Resour. 4:359–361.

Elmer, K. R., T. K. Lehtonen, and A. Meyer. 2009. Color assortative mating contributes to sympatric divergence of neotropical cichlid fish. Evolution 63:2750–2757.

Etges, W. J., and D. A. Tripodi. 2008. Premating isolation is determined by larval rearing substrates in cactophilic *Drosophila mojavensis*. VIII. Mating success mediated by epicuticular hydrocarbons within and between isolated populations. J. Evol. Biol. 21:1641–1652.

Evanno, G., S. Regnaut, and J. Goudet. 2005. Detecting the number of clusters of individuals using the software STRUCTURE: a simulation study. Mol. Ecol. 14:2611–2620.

Felsenstein, J. 1981. Skepticism towards Santa Rosalia, or why are there so few kinds of animals? Evolution 35:124–138.

Fisher, H. S., B. B. M. Wong, and G. G. Rosenthal. 2006. Alteration of the chemical environment disrupts communication in a freshwater fish. Proc. R. Soc. B. 273:1187–1193.

Gilman, R. T., and G. M. Kozak. 2015. Learning to speciate: the biased learning of mate preferences promotes adaptive radiation. Evolution 69:3004–3012.

Grant, P. R., and B. R. Grant. 1997. Hybridization, sexual imprinting, and mate choice. Am. Nat. 149:1–28.

Hall, E. R. 1981. The mammals of North America. Second edition. John Wiley & Sons, New York.

Hall, E. R., and K. R. Kelson. 1959. The mammals of North America. Ronald Press Company, New York.

Hartung, T. G., and D. A. Dewsbury. 1979. Paternal behavior in six species of muroid rodents. Behav. Neural Biol. 26:466–478.

Howell, A. H. 1921. A biological survey of Alabama. North Am. Fauna 45:1–89.

Immelmann, K. 1975. Ecological significance of imprinting and early learning. Annu. Rev. Ecol. Evol. Syst. 6:15–37.

Irwin, D. E., and T. Price. 1999. Sexual imprinting, learning and speciation. Heredity 82:347–354.

Jakobsson, M., and N. a Rosenberg. 2007. CLUMPP: a cluster matching and permutation program for dealing with label switching and multimodality in analysis of population structure. Bioinformatics 23:1801–1806.

Kendrick, K. M., M. R. Hinton, K. Atkins, M. A. Haupt, and J. D. Skinner. 1998. Mothers determine sexual preferences. Nature 395:229–230.

Kopp, M., R. J. Safran, M. R. Servedio, R. L. Rodr, T. C. Mendelson, M. E. Hauber, E. C. Scordato, C. N. Balakrishnan, L. B. Symes, D. M. Zonana, and G. S. Van Doorn. n.d. Mechanisms of assortative mating in speciation with gene flow: connecting theory and empirical research. Am. Nat. in press.

Kozak, G. M., and J. W. Boughman. 2009. Learned conspecific mate preference in a species pair of sticklebacks. Behav. Ecol. 20:1282–1288.

Kozak, G. M., M. L. Head, and J. W. Boughman. 2011. Sexual imprinting on ecologically divergent traits leads to sexual isolation in sticklebacks. Proc. R. Soc. B. 278:2604–2610.

Lackey, J. A., D. G. Huckaby, and B. G. Ormiston. 1985. Peromyscus leucopus. Mamm. Species 247:1–10.

Laland, K. N. 1994. On the evolutionary consequences of sexual imprinting. Evolution 48:477–489.

Latour, Y., M. Perriat-Sanguinet, P. Caminade, P. Boursot, C. M. Smadja, and G. Ganem. 2014. Sexual selection against natural hybrids may contribute to reinforcement in a house mouse hybrid zone. Proc. R. Soc. B. 281:20132733.

Lovecky, D. V., D. Q. Estep, and D. A. Dewsbury. 1979. Copulatory behavior of cotton mice (*Peromyscus gossypinus*) and their reciprocal hybrids with white-footed mice (*P. leucopus*). Anim. Behav. 27:371–375.

Lunter, G., and M. Goodson. 2011. Stampy: a statistical algorithm for sensitive and fast mapping of Illumina sequence reads. Genome Res. 21:936–939.

Maddock, M. B., and W. D. Dawson. 1974. Artificial insemination of deermice (*Peromyscus maniculatus*) with sperm from other rodent species. J. Embryol. Exp. Morphol. 31:621–634.

Matsubayashi, K. W., and H. Katakura. 2009. Contribution of multiple isolating barriers to reproductive isolation between a pair of phytophagous ladybird beetles. Evolution 63:2563–2580.

McCarley, W. H. 1954a. Natural hybridization in the *Peromyscus leucopus* species group of mice. Evolution 8:314–323.

McCarley, W. H. 1954b. The ecological distribution of the *Peromyscus leucopus* species group in eastern Texas. Ecology 35:375–379.

McCarley, W. H. 1954c. Fluctuations and structure of *Peromyscus gossypinus* populations in eastern Texas. J. Mammal. 35:526–532.

McCarley, W. H. 1963. Distributional relationships of sympatric populations of *Peromyscus leucopus* and *P. gossypinus*. Ecology 44:784–788.

Mcvean, G. 2009. A genealogical interpretation of principal components analysis. PLoS Genet. 5:e1000686.

McCarty, R., and C. H. Southwick. 1977a. Cross-species fostering: effects on the olfactory preference of *Onychomys torridus* and *Peromyscus leucopus*. Behav. Biol. 19:255–260.

McCarty, R., and C. H. Southwick. 1977b. Patterns of parental care in two Cricetid rodents, *Onychomys torridus* and *Peromyscus leucopus*. Anim. Behav. 25:945–948.

McKenna, A., M. Hanna, E. Banks, A. Sivachenko, K. Cibulskis, A. Kernytsky, K. Garimella, D. Altshuler, S. Gabriel, M. Daly, and M. A. DePristo. 2010. The Genome Analysis Toolkit: a MapReduce framework for analyzing next-generation DNA sequencing data. Genome Res. 20:1297–1303.

Mendelson, T. C. 2003. Sexual isolation evolves faster than hybrid inviability in a diverse and sexually dimorphic genus of fish (Percidae: *Etheostoma*). Evolution 57:317–327.

Montero, I., M. Teschke, and D. Tautz. 2013. Paternal imprinting of mating preferences between natural populations of house mice (*Mus musculus domesticus*). Mol. Ecol. 22:2549–2562.

Nei, M. 1972. Genetic distance between populations. Am. Nat. 106:283–292.

Noor, M. A. F. 1997. How often does sympatry affect sexual isolation in *Drosophila*? Am. Nat. 149:1156–1163.

Nosil, P. 2007. Divergent host plant adaptation and reproductive isolation between ecotypes of *Timema cristinae* walking sticks. Am. Nat. 169:151–162.

Nosil, P., R. Riesch, and M. Muschick. 2013. Climate affects geographic variation in host-plant but not mating preferences of *Timema cristinae* stick-insect populations. Evol. Ecol. Res. 15:1–16.

Patterson, N., A. L. Price, and D. Reich. 2006. Population structure and eigenanalysis. PLoS Genet. 2:2074–2093.

Peterson, B. K., J. N. Weber, E. H. Kay, H. S. Fisher, and H. E. Hoekstra. 2012. Double digest RADseq: an inexpensive method for *de novo* SNP discovery and genotyping in model and non-model species. PLoS One 7:e37135.

Platt, R. N., B. R. Amman, M. S. Keith, C. W. Thompson, and R. D. Bradley. 2015. What Is Peromyscus? Evidence from nuclear and mitochondrial DNA sequences suggests the need for a new classification. J. Mammal. 96:708–719.

Pournelle, G. H. 1952. Reproduction and early post-natal development of the cotton mouse, *Peromyscus gossypinus*. J. Mammal. 33:1–20.

Price, P. K., and M. L. Kennedy. 1980. Genic relationships in the white-footed mouse, *Peromyscus leucopus*, and the cotton mouse, *Peromyscus gossypinus*. Am. Midl. Nat. 103:73–82.

Pritchard, J. K., M. Stephens, and P. Donnelly. 2000. Inference of population structure using multilocus genotype data. Genetics 155:945–959.

Ramsey, J., H. D. Bradshaw, and D. W. Schemske. 2003. Components of reproductive isolation between the monkeyflowers *Mimulus lewisii* and *M. cardinalis* (Phrymaceae). Evolution 57:1520–1534.

Robbins, L. W., M. H. Smith, M. C. Wooten, and R. K. Selander. 1985. Biochemical polymorphism and its relationship to chromosomal and morphological variation in *Peromyscus leucopus* and *Peromyscus gossypinus*. J. Mammal. 66:498–510.

Roehrs, Z. P., J. B. Lack, C. E. S. Jr., C. J. Seiden, R. Bastarache, W. D. Arbour, D. M. L. Jr., and R. A. Van Den Bussche. 2012. Mammals of Red Slough Wildlife Management Area, with comments on McCurtain county, Oklahoma. Texas Tech Univ. Occas. Pap. 309:1–24.

Rolán-Alvarez, E., and A. Caballero. 2000. Estimating sexual selection and sexual isolation effects from mating frequencies. Evolution 54:30–36.

Rosenberg, N. A. 2004. Distruct: a program for the graphical display of population structure. Mol. Ecol. Notes 4:137–138.

Rosenthal, G. G. 2017. Mate choice: the evolution of sexual decision making from microbes to humans. Princeton University Press, Princeton.

Schug, M. D., S. H. Vessey, and E. M. Underwood. 1992. Paternal behavior in a natural population of mice (*Peromyscus leucopus*). Am. Midl. Nat. 127:373–380.

Seehausen, O., J. M. van A. Jacques, and F. Witte. 1997. Cichlid fish diversity threatened by eutrophication that curbs sexual selection. Science 277:1808–1811.

Servedio, M. R., S. A. Sæther, and G.-P. Sætre. 2009. Reinforcement and learning. Evol. Ecol. 23:109–123.

Slagsvold, T., B. T. Hansen, L. E. Johannessen, and J. T. Lifjeld. 2002. Mate choice and imprinting in birds studied by cross-fostering in the wild. Proc. R. Soc. B. 269:1449–1455.

Smadja, C. M., and R. K. Butlin. 2011. A framework for comparing processes of speciation in the presence of gene flow. Mol. Ecol. 20:5123–5140.

Soltis, D. E., A. B. Morris, J. S. McLachlan, P. S. Manos, and P. S. Soltis. 2006. Comparative phylogeography of unglaciated eastern North America. Mol. Ecol. 15:4261–4293.

Taylor, R. J., and H. McCarley. 1963. Vertical distribution of *Peromyscus leucopus* and *P. gossypinus* under experimental conditions. Southwest. Assoc. Nat. 8:107–115.

ten Cate, C., and G. Mug. 1984. The development of mate choice in zebra finch females. Behaviour 90:125–150.

ten Cate, C., and C. Rowe. 2007. Biases in signal evolution: learning makes a difference. Trends Ecol. Evol. 22:380–387.

ten Cate, C., M. N. Verzijden, and E. Etman. 2006. Sexual imprinting can induce sexual preferences for exaggerated parental traits. Curr. Biol. 16:1128–1132.

ten Cate, C., and D. R. Vos. 1999. Sexual imprinting and evolutionary processes in birds: a reassessment. Pp. 1–31 in P. J. Slater, J. S. Rosenblatt, C. T. Snowdon, and T. J. Roper, eds. Mate Choice. Academic Press.

Tramm, N. A., and M. R. Servedio. 2008. Evolution of mate-choice imprinting: competing strategies. Evolution 62:1991–2003.

Turner, L. M., D. J. Schwahn, and B. Harr. 2011. Reduced male fertility is common but highly variable in form and severity in a natural mouse hybrid zone. Evolution 66:443–458.

Verzijden, M. N., and C. ten Cate. 2007. Early learning influences species assortative mating preferences in Lake Victoria cichlid fish. Biol. Lett. 3:134–136.

Verzijden, M. N., and G. G. Rosenthal. 2011. Effects of sensory modality on learned mate preferences in female swordtails. Anim. Behav. 82:557–562.

Verzijden, M. N., R. F. Lachlan, and M. R. Servedio. 2005. Female mate-choice behavior and sympatric speciation. Evolution 59:2097–2108.

Verzijden, M. N., C. Cate, M. R. Servedio, G. M. Kozak, J. W. Boughman, and E. I. Svensson. 2012a. The impact of learning on sexual selection and speciation. Trends Ecol. Evol. 27:511–519.

Verzijden, M. N., Z. W. Culumber, and G. G. Rosenthal. 2012b. Opposite effects of learning cause asymmetric mate preferences in hybridizing species. Behav. Ecol. 23:1133–1139.

Vos, D. R. 1995. Sexual imprinting in zebra-finch females: do females develop a preference for males that look like their father? Ethology 99:252–262.

Weber, J. N., M. B. Peters, O. V Tsyusko, C. R. Linnen, C. Hagen, N. A. Schable, T. D. Tuberville, A. M. McKee, S. L. Lance, K. L. Jones, H. S. Fisher, M. J. Dewey, H. E. Hoekstra, and T. C. Glenn. 2010. Five hundred microsatellite loci for *Peromyscus*. Conserv. Genet. 11:1243–1246.

Wilson, R. C., T. Vacek, D. L. Lanier, and D. A. Dewsbury. 1976. Open-field behavior in muroid rodents. Behav. Biol. 506:495–506.

Wolfe, J. L., and A. V Linzey. 1977. Peromyscus gossypinus. Mamm. Species 70:1–5.

Yukilevich, R., and J. R. True. 2008. Incipient sexual isolation among cosmopolitan *Drosophila melanogaster* populations. Evolution 62:2112–2121.

Zimmerman, E. G., C. W. Kilpatrick, and B. J. Hart. 1978. The genetics of speciation in the rodent genus *Peromyscus*. Evolution 32:565–579.

